# Membrane Sensing Peptides -Enhanced SiMoA Platform for the Detection of HER2 on Extracellular Vesicles in Metastatic Breast Cancer Patients

**DOI:** 10.1101/2025.09.18.676908

**Authors:** Arianna Bonizzi, Paola Gagni, Francesca Piccotti, Marta Truffi, Sara Albasini, Serena Mazzucchelli, Lorena Signati, Tiziana Triulzi, Elda Tagliabue, Alessandro Gori, Giorgia Esposito, Francesca Gorgoglione, Ilaria Tagliolini, Fabio Corsi, Carlo Morasso

## Abstract

Extracellular vesicles (EVs) offer a promising avenue for non-invasive, real-time monitoring of metastatic breast cancer (mBC), but clinical application as a liquid biopsy is hindered by their heterogeneity and low abundance. Here we present a Single Molecule Array (SiMoA) platform enhanced by membrane sensing peptides (MSP) for the highly sensitive detection of HER2 on EV membranes (EVs-HER2) and general EVs population (CD9+) directly from plasma samples of mBC patients. The MSP-based SiMoA assay demonstrated superior sensitivity and specificity compared to conventional antibody-based assays, allowing the detection of lower amounts of EVs and discriminating EVs derived from breast cancer patient-derived organoids (BC-PDO) from healthy control-derived organoids (HC-PDO).

Concerning the analysis of EVs in plasma samples (n=49 mBC patients, n=30 healthy controls), we observed significantly lower CD9+ EVs levels in mBC patients relative to healthy controls, a trend consistently confirmed across assays. Notably, EVs-HER2 levels were significantly enriched in HER2-positive patients and correlated with clinical HER2 status assessed by immunohistochemistry. Besides, lower CD9+ EVs levels were associated with poorer clinical outcomes, highlighting the potential prognostic utility of EV quantification.

Our findings underscore the potential of MSP-enhanced SiMoA platforms for accurate, minimally invasive monitoring of EVs-HER2 in mBC and for monitoring CD9+ EVs levels to assess disease progression.

## 1. INTRODUCTION

HER2-positive breast cancer (HER2+ BC) accounts for 15-20% of cases and is associated with aggressive progression and higher recurrence risk^1^. Current therapeutic strategies for HER2+ BC primarily involve antibodies targeting specific HER2 epitopes^2^. However, monitoring HER2 status during treatment remains challenging in metastatic patients, as biopsies are often not feasible, and HER2 assessments is often performed in earlier treatment phases and may no longer accurately reflect the disease’s continuously evolving molecular profile^3–5^.

Liquid biopsy using extracellular vesicles (EVs) potentially offers non-invasive, real-time molecular assessment and monitoring of metastatic breast cancer^6,7^. Still, despite more than ten years of research in the field, the clinical implementation of EVs as biomarker remains limited due to complex isolation and analysis protocols^8^. Furthermore, EVs in blood are highly heterogenous as originate from multiple organs and tissues and may undergo complex rearrangements through the interaction with circulating factors—a phenomenon known as the “extracellular vesicle corona” ^9,10^.

To detect cancer released EVs in a clinically useful way there is thus the pressing need of new bioanalytical assays able to detect extremely low amount of target in a way that is scalable for the study of multiple groups of subjects in a short time^11^.

In this context, Single Molecule Array (SiMoA) technology has emerged as a powerful method that integrates the magnetic immunoseparation of EVs with an advanced digital quantification, analysing binding events individually on each magnetic bead. According to the most recent MISEV guidelines^12^, immunoseparation is the approach able to provide purest EVs, though typically with lower recovery rates^13^. Its combination with the digital quantification of markers, unlock the possibility to detect quickly ultra-low concentrations of EVs in unprocessed plasma with high sensitivity^14^.

SiMoA has previously demonstrated success in the neurological field for similarly challenging applications, such as detecting proteins originating from the central nervous system as neurofilament light chain (NfL), and Glial fibrillary acidic protein (GFAP) directly in blood at ultra-low concentrations^15–18^. Our group previously demonstrated the successful application of SiMoA for rapid and sensitive EVs detection in the plasa of early breast cancer patients^19,20^. To further extend the SiMoA analysis at the study of EVs not necessarily presenting tetraspanins as markers on their membrane^21^, we also previously proposed the use of a membrane sensing peptides (MSP) as universal EV-binding probes enabling integrated and unbiased isolation and phenotyping^22^.

Building upon these previous findings, the present study aims to demonstrate the practical feasibility and robustness of the SiMoA MSP-based assay as method to evaluate the clinical relevance of HER2 on EV membranes (EVs-HER2) and general EVs (CD9+) levels in metastatic breast cancer (mBC) patients.

To provide specificity, the assay utilizes an anti-HER2 antibody targeting the extracellular domain of HER2, on an epitope distinct from those targeted by current anti-HER2 therapies, such as Trastuzumab and Pertuzumab^23,24^, for the quantification of HER2 expressed on the surface of EVs Specifically, the anti-HER2 antibody used in the assays is a monoclonal antibody, designated MGR2 (IgG1), obtained by immunizing mice with live carcinoma cells overexpressing the HER2 oncoprotein. The epitope defined by MGR2 is discontinuous and hydrophobic interactions are involved in binding with HER2 oncoprotein indicating the ability of this antibody to recognize HER2 in the conformation it has in tumours cells^25^. Levels of plasmatic general circulating EVs were also measured using the MSP conjugated beads paired with and anti-CD9 monoclonal antibody. A second set of assays employing magnetic beads using a conventional anti-CD63 antibody capture strategy was also developed and the two approach were comparatively evaluated. The four developed SiMoA assays were first tested on EVs isolated from the culture medium of breast cancer patient-derived organoids (PDOs). After validating their analytical performance, these assays were employed for the direct quantification of EVs in unprocessed plasma samples from 49 mBC patients and 30 healthy controls (HC). This analysis enabled the correlation between HER2 levels on EVs and the clinical HER2 status assessed through standard immunohistochemical (IHC). Additionally, the influence of pre-analytical factors such as haemolysis and lipemia, and other haematological parameters on EVs and EVs-HER2 levels was also evaluated.

## 2. RESULTS

### 2.1 Characterization of EVs isolated from PDO culture media

EVs were isolated from culture media of a breast cancer derived PDO (BC-PDO) using SEC. The PDO was established from a primary BC biopsy sample, and cultured according to a previously published protocol^26^. Although the original tissue biopsy did not show HER2 overexpression (histopathological scored as 1+), the IHC analysis of the cultured PDO revealed evidences of a weak HER2 expression (Figure 1A).

**Figure 1.**
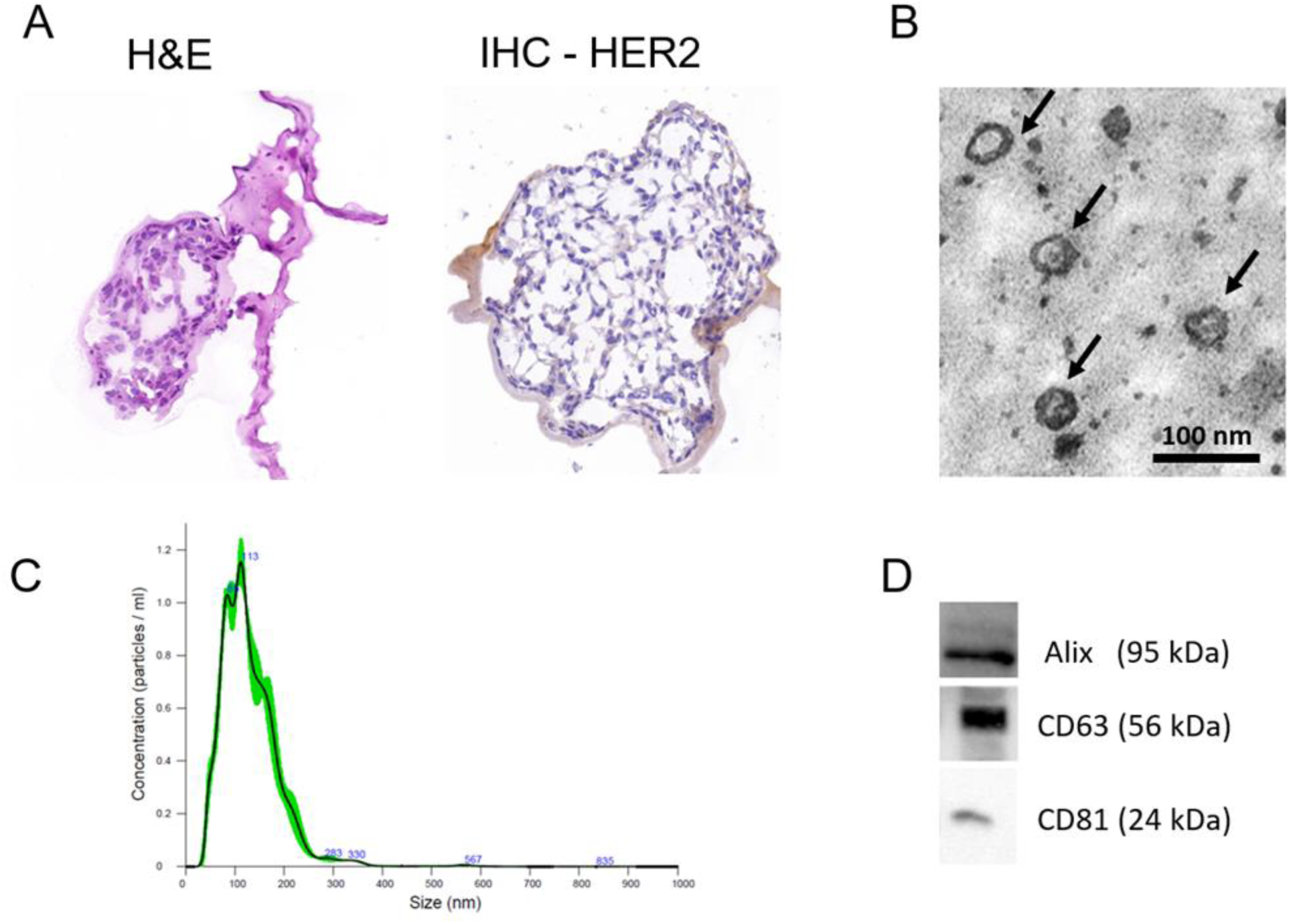
A) Haematoxylin and eosin (H&E) staining and HER2 IHC analysis of BC-PDO; B) Representative TEM images of the EVs. Scalebar = 100 nm; C) NTA analysis of EVs; D) Western blot analysis of EVs markers (Alix, CD63, and CD81).

Isolated EVs were characterized following the MISEV guidelines^12,27^. TEM imaging verified the typical spherical morphology and membrane integrity, confirming an average vesicle size around 100 nm (Figure 1B). NTA was used to characterize the size distribution of the EVs preparations, revealing a distribution of two prominent peaks at approximately 85 nm and 113 nm, reflecting EVs heterogeneity typical of cancer-derived samples (Figure 1C). WB analysis confirmed the presence of typical EVs markers Alix, CD63 and CD81 validating EVs (Figure 1D). Collectively, these data confirm successful collection of EVs from the culture media.

### 2.2 Characterization of the SiMoA assays

Two SiMoA assay platforms using anti-CD63 or MSP-conjugated magnetic beads were comparatively evaluated for their sensitivity in detecting EVs. General EVs detection was performed using the beads coupled with a commercially available biotinylated anti-CD9 monoclonal antibody, while for detecting EVs-HER2, beads were coupled with the biotinylated anti-HER2 monoclonal antibody MGR2, which specifically targets the extracellular domain of HER2^24^. Increasing amounts of PDO-derived EVs measured as total protein content using the BCA assay, were loaded per well for each assay. The Average Enzyme per Bead Activity (AEB) signals obtained for both assays were analysed using a four-parameter logistic (4PL) regression model to estimate performance (Figure 2)^28,29^.

**Figure 2.**
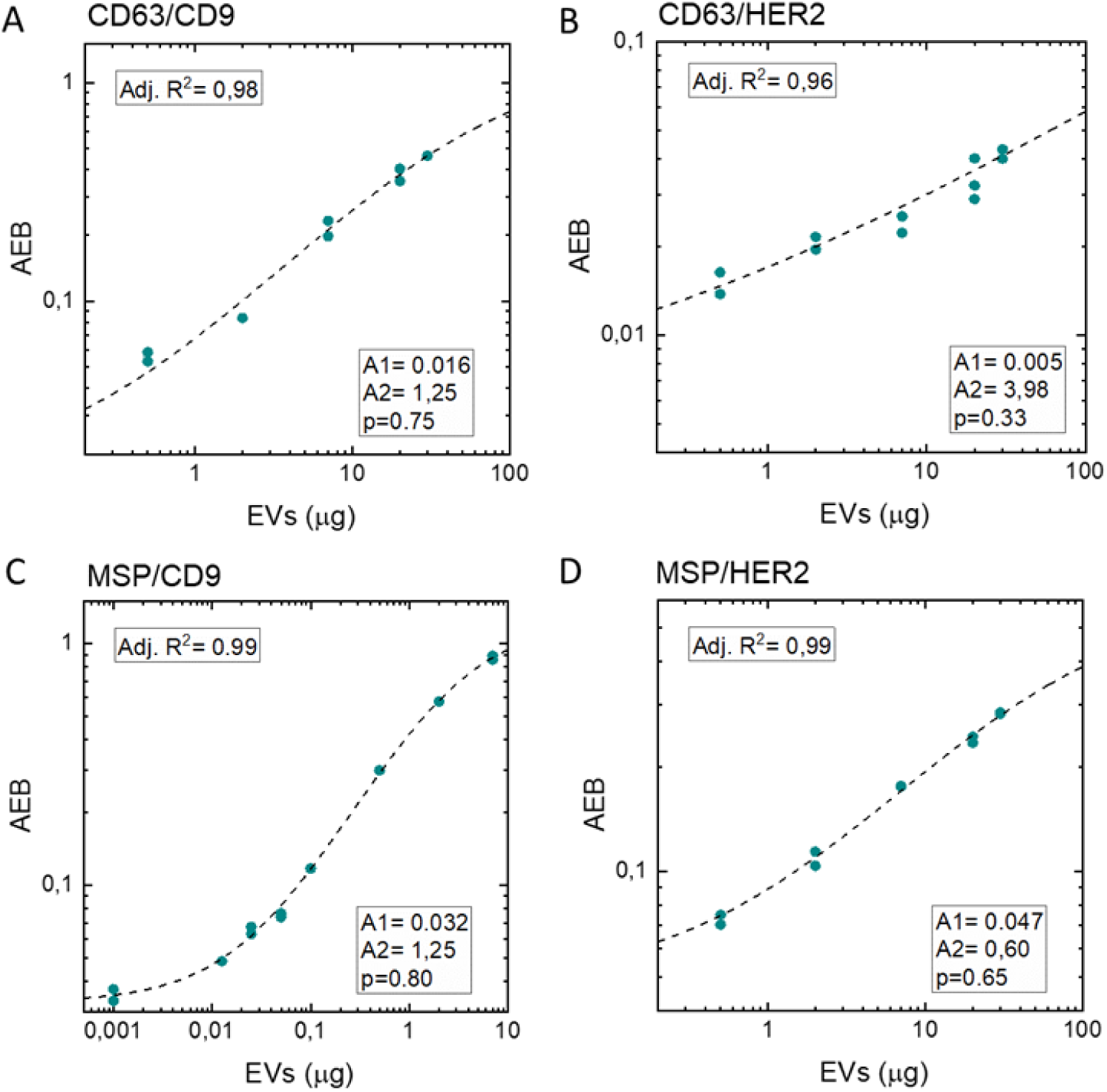
Response curve of the assays at different amount of EVs derived from the BC-PDO. A) CD63/CD9 assay. B) CD63/HER2 assay. C) MSP/CD9 assay. D) MSP/HER2 assay. Data were fitted with a four-parameter logistic (4PL) curve. A1: Minimum asymptote. A2: Maximum asymptote. p: Hill’s slope.

The comparative results demonstrated a clear dose-response relationship for all assays analysed, reflected by consistently high adjusted R² values across all calibration curves (Figure 2). MSP-based assays (MSP/CD9 and MSP/HER2) exhibited higher baseline signals (Minimum asymptote (A1) = 0.032 for MSP/CD9, A1 = 0.047 for MSP/HER2) compared to their CD63-based counterparts (Minimum asymptote (A1) = 0.016 for CD63/CD9, A1 = 0.005 for CD63/HER2), indicative of higher background noise levels associated with the MSP beads. Nevertheless, the MSP-based assays also demonstrated significantly enhanced sensitivity and superior overall analytical performance, as evidenced by the notably higher Hill’s slope (p) derived from the regression analysis (p = 0.65 for MSP/HER2 versus 0.33 for CD63/HER2). Hill slope (p) is the exponential term in the 4PL fit that controls the steepness of the relationship between signal and analyte concentration, indicating the strength of this dependence (Figure S2). Furthermore, MSP beads allowed the detection of substantially lower EVs concentrations, prompting the recalibration of assay ranges toward lower values and thus extending the effective dynamic range of quantification.

Overall, the SiMoA assays developed operate in the digital detection mode, where AEB is directly derived from the fraction of enzymatically active beads (fON), remained optimal, as indicated by AEB values consistently below 1 across all assays^30^.

To further evaluate the response of the two developed assays for EVs-HER2 detection, EVs were also isolated from the culture media of a second PDO derived from healthy breast tissue, which does not exhibit HER2 receptor expression on the membrane at IHC (Figure S1). Equal protein amounts (20 µg) of PDO-derived EVs from BC-PDO and HC-PDO were analysed. Results presented in Figure 3 show that the MSP-based assay produced higher AEB signals compared to the CD63-based assay. Notably, the CD63/HER2 assay did not detect significant differences between the EVs populations derived from cancerous and healthy PDOs, confirming the low sensitivity of the CD63 based assay for the detection of EVs-HER2 (Figure 3A). Conversely, the MSP-based assay demonstrated better performances, clearly distinguishing the cancer-derived EVs due to their significantly higher HER2 levels (Figure 3B). These findings reinforce the superior response of the MSP-based assay for precise quantification of EVs-HER2. We speculate this result may reflect the loss of HER2+ EVs subpopulations when CD63 is used as capturing probes. MSP, acting as pan-vesicular probes, is able to restore the information as it targets a broader population of EVs. Of note, this phenomenon was previously observed for EVs markers in cardiovascular risk stratification^22^, and this is consistent with the growing evidence that tetraspanin-responsive EVs represent only a partial fraction of circulating EVs.+

**Figure 3.**
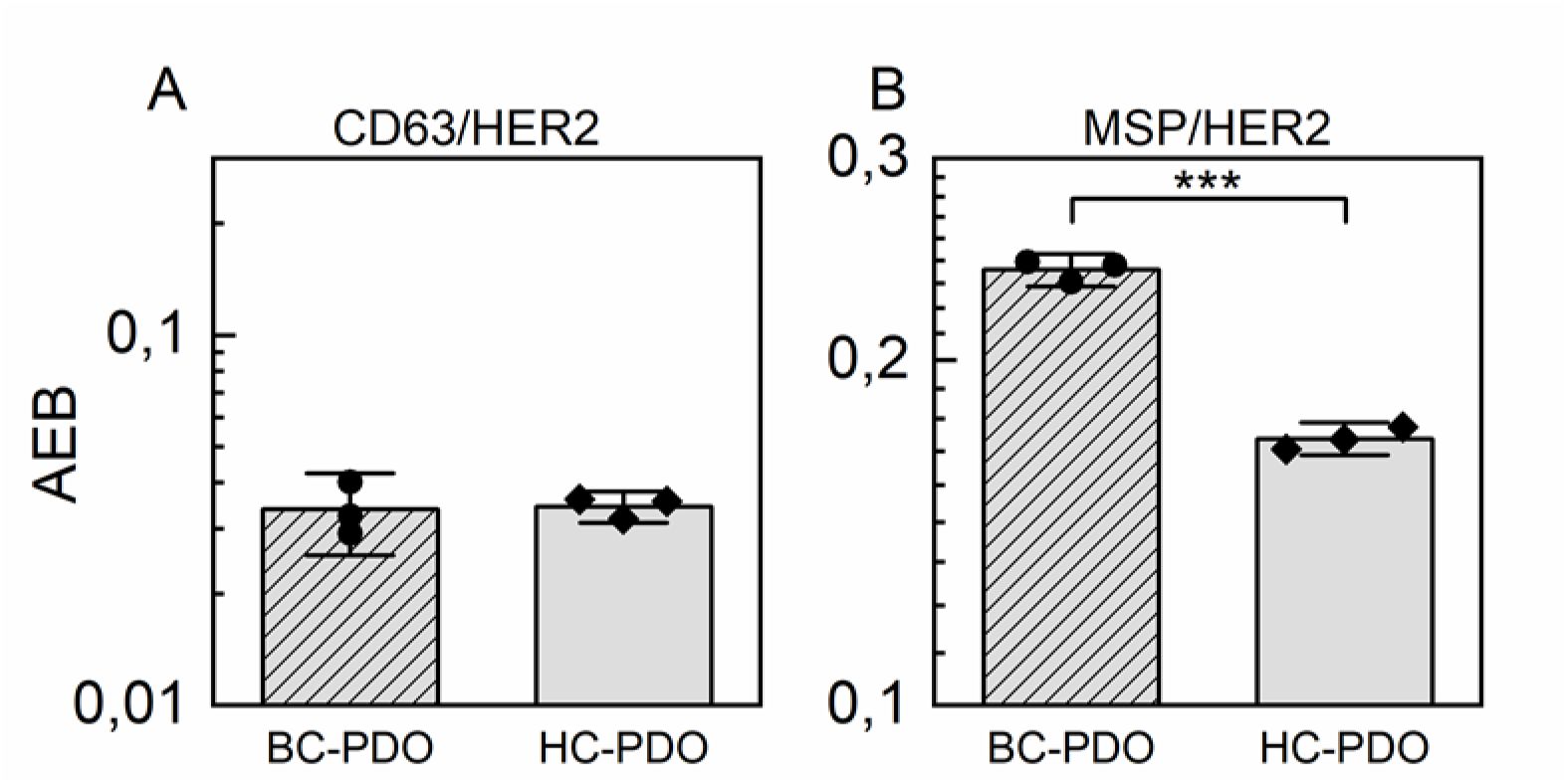
EVs-HER2 detection from BC-PDO and HC-PDO derived EVs by two capture strategies: A) CD63/HER2 assay. B) MSP/HER2 assay. p-value ***<0,0005.

Collectively, our results suggest that while both assay configurations are suitable for quantifying EVs levels and specifically HER2-expressing EVs, the two approaches have different characteristics and the MSP-based SiMoA methodology presents a better response to the presence of EVs.

### 2.3 Analysis of blood plasma samples

We thus extended the analysis to plasma samples from mBC patients and HC. Specifically, 17 HER2-positive (HER2+) and 32 HER2-negative (HER2-) mBC patients, along with 30 HC, were enrolled. Plasma was collected and analysed without further processing with the two sets of SiMoA assays, enabling the direct detection of general EVs (via anti-CD9) and HER2-expressing EVs (via anti-HER2). Since it was not possible analyse all samples with all assays due to limits in the amount of plasma samples available, in some cases the number of patients considered is lower than the total as reported in Table 1.

**Table 1:** Correlation (Spearman) between assays, age, BMI and preanalytical factors.

| Assay: | Unit | n° | CD63/CD9<br>*R*<br>(*p*) | MSP/CD9<br>*R*<br>(*p*) | CD63/HER2<br>*R*<br>(*p*) | MSP/HER2<br>*R*<br>(*p*) |
| --- | --- | --- | --- | --- | --- | --- |
| CD63/CD9 | AEB | mBC n = 48; HC<br>30 |  |  |  |  |
| MSP/CD9 | AEB | mBC n = 48; HC<br>26 | 0,16<br>(0,16) |  |  |  |
| CD63/HER2 | AEB | mBC n = 45; HC<br>30 | **0,32**<br>**(0,005)** | 0,06<br>(0,60) |  |  |
| MSP/HER2 | AEB | mBC n = 49; HC<br>30 | **0,36**<br>**(0,0012)** | **0,54**<br>**(<0.0001)** | **0,27**<br>**(0,017)** |  |
| Age | Years | mBC n = 49; HC<br>30 | -0,06<br>(0,59) | -0,07<br>(0,53) | 0,02<br>(0,82) | -0,01<br>(0,91) |
| BMI | Kg/m<sup>2</sup> | mBC n = 46; HC<br>6 | -0,11<br>(0,44) | -0,21<br>(0,14) | -0,21<br>(0,15) | **-0,33**<br>**(0,016)** |
| Hemolysis | 414-385 nm | mBC n = 49; HC<br>30 | 0,21<br>(0,07) | 0,20<br>(0,08) | -0,01<br>(0,94) | 0,19<br>(0,09) |
| Lipemia | 780 nm | mBC n = 49; HC<br>30 | -0,04<br>(0,69) | -0,18<br>(0,13) | -0,01<br>(0,94) | -0,15<br>(0,17) |

Values in Bold show statistical significance ( $p < 0.05$ )

Correlation analyses were performed to assess the consistency between the two assay platforms for quantifying CD9+ EVs and EVs-HER2. For circulating CD9+ EVs, the results obtained using the CD63 and MSP bead assays showed no significant correlation (*R* = 0,16) suggesting that the two assays target different subpopulations of circulating EVs. Instead, a weak correlation was found between the assays for EVs-HER2 quantification (R = 0,27), as reported in Table 2. A moderate correlation was reported between EVs-HER2 and the CD9+ EVs measured both with MSP assays (R = 0,54), a result that confirm the fact that the HER2 measured in this assay is effectively transported by EVs.

**Table 2.** Characteristics of the enrolled cohorts.

| Parameters | HC (n=30) | mBC (n=49) | *p* |
| --- | --- | --- | --- |
| Age (years) | 55,67 ± 14,14 | 62,93 ± 13,64 | * |
| BMI (Kg/m<sup>2</sup>) | 23,84 ± 5,70 | 26,04 ± 7,36 | n.s |
| HER2 + |  | 17 |  |
| IHC 3+ |  | 7 |  |
| IHC 2+ FISH (+) |  | 10 |  |
| HER2 – |  | 32 |  |
| IHC 2+ FISH (-) |  | 9 |  |
| IHC 1+ |  | 14 |  |
| IHC 0+ |  | 9 |  |

In accordance with MISEV guidelines and the document drafted by the ISEV Blood EV Task Force (MIBlood-EV)^31^, we evaluated the influence of pre-analytical factors on our data. No significant effect of haemolysis and lipemia was observed (Table 1). Age and BMI were also checked. While no effect of age was found, a low inverse correlation was found between BMI and EVs-HER2 measured using MSP beads. Additionally, for mBC samples, we evaluated the impact of other routinely measured variables—complete blood count parameters for platelets, erythrocytes, and leukocytes— and the serum concentrations of the divalent electrolytes magnesium and calcium whose presence could in principle affect the interaction between beads and EVs. No significant correlation was observed between the SiMoA assays and other haematological values (Table S1).

### 2.4 EVs and EV-HER2 in metastatic breast cancer

Plasma levels of general EVs (detected using anti-CD9) and EVs-HER2 (detected using anti-HER2) were compared between mBC patients (n=49) and HC (n=30) using the two different capture strategies.

Rather surprisingly, our data revealed that CD9+ EVs were significantly lower in mBC compared to HC across both assay platforms (Figure 4A and 4B). Although EVs-HER2 could be detected at very low concentrations in patient, the CD63/HER2 assay did not reveal a significant difference between mBC and HC (Figure 4C). In contrast, the MSP-based assay for EVs-HER2 detection showed a clear statistically significant difference between the two groups (Figure 4D) confirming the fact that mBC are characterized by lower levels of both EVs and EVs-HER2.

**Figure 4:**
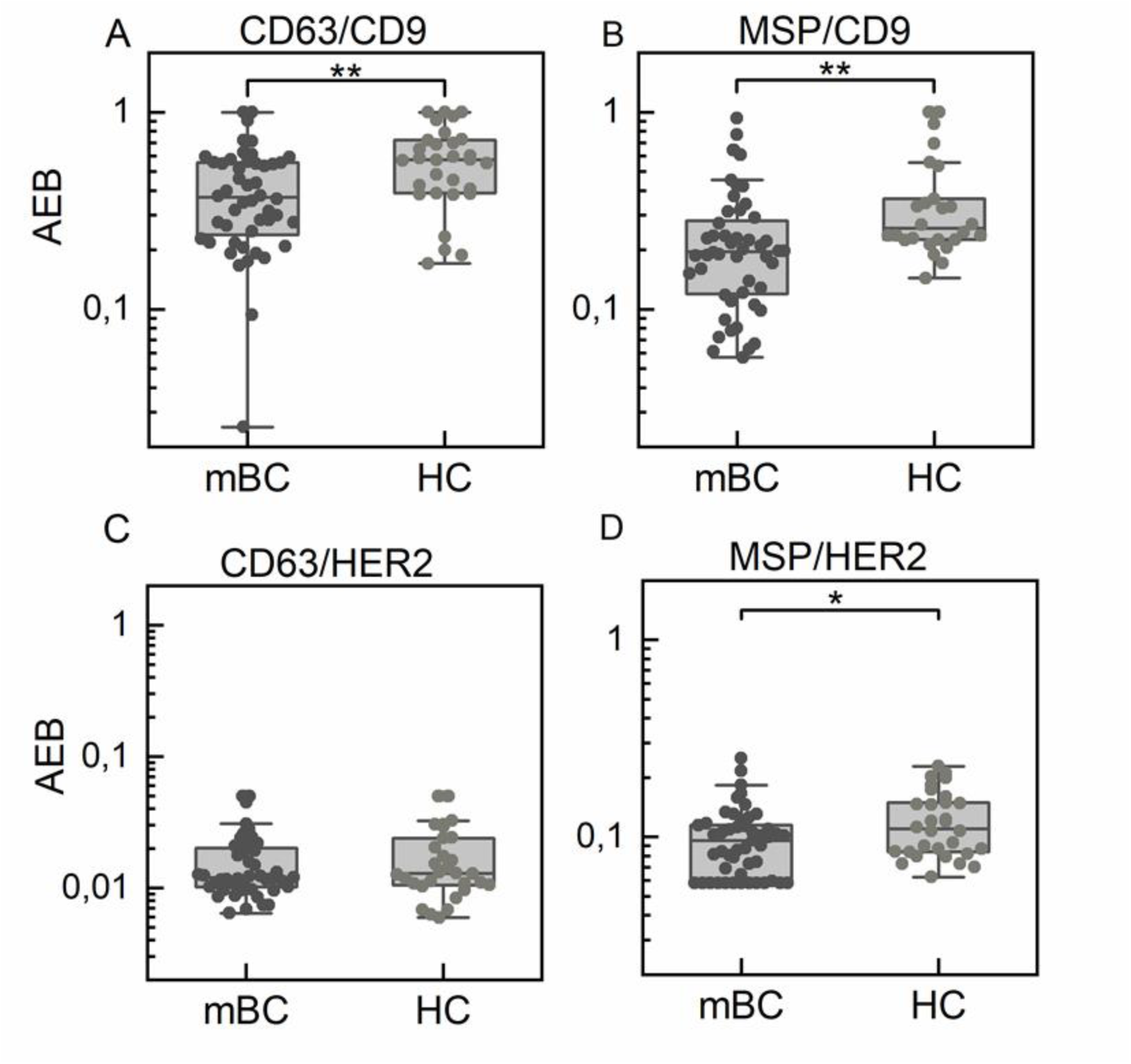
Plasma levels of CD9+ EVs (A, B) and EVs-HER2 (C, D) measured in mBC and HC. Data are shown as box and whisker plots. Each data point represents an individual subject analysed. p-value *<0,05; **<0,005.

When mBC patients were stratified between HER2+ (n=17) and HER2-(n=32) subgroups, neither assay demonstrated significant differences in circulating CD9+ EVs levels between the two groups of patients (Figure 5A and 5B). Both assays consistently indicated that EVs-HER2 levels were significantly enriched in HER2+ mBC patients compared to HER2-patients, in line with our expectations (Figure 5C and 5D).

**Figure 5:**
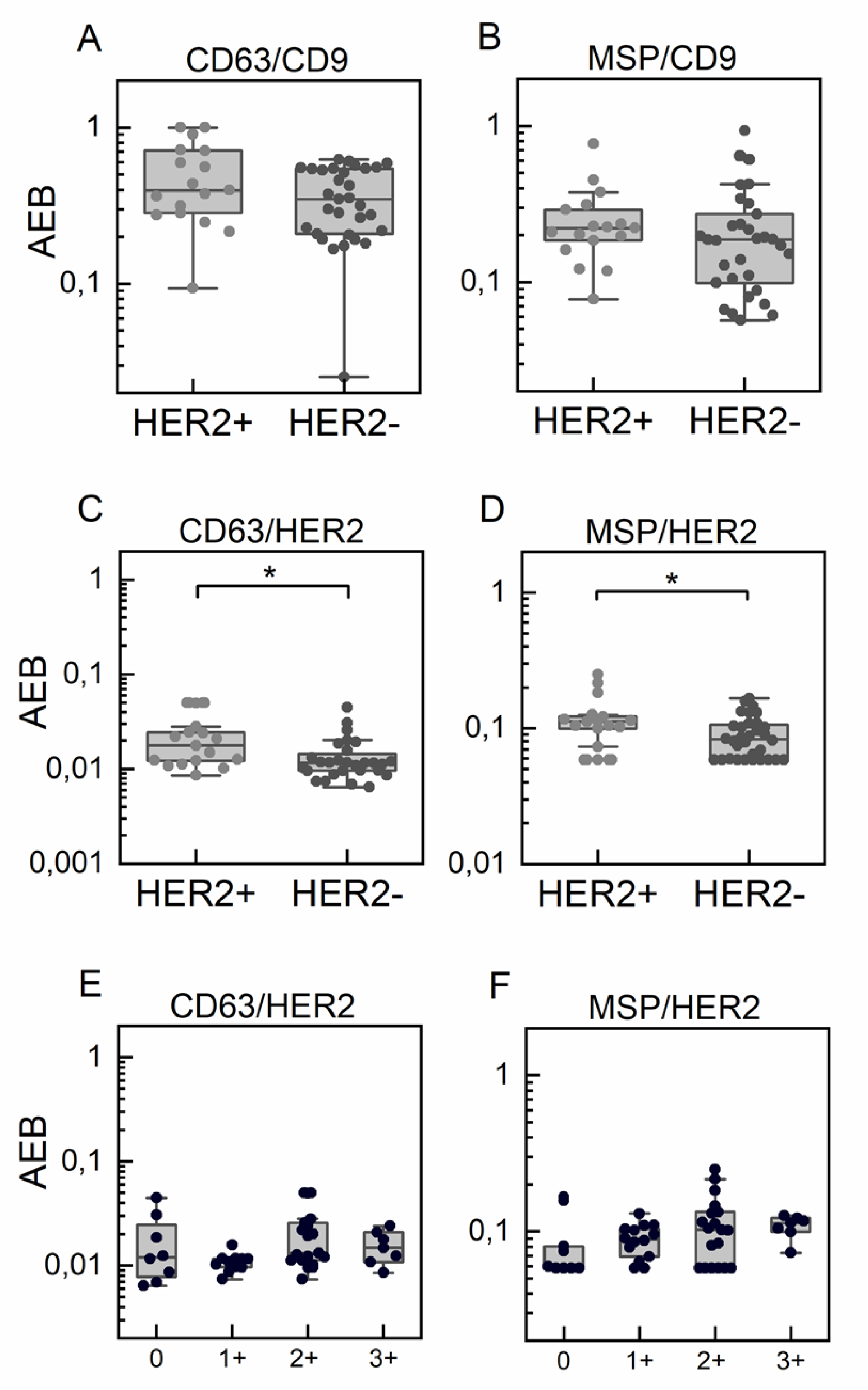
A, B) Plasma levels of EVs measured using the CD63/CD9 assay (HER2+ n = 17; HER2-n = 31) and with the MSP/CD9 assay (HER2+ n = 17; HER2-n = 31). C, D) Plasma levels of EVs-HER2 measured using the CD63/HER2 assay (HER2+ n = 17; HER2-n = 28) and with the MSP/HER2 assay (HER2+ n = 17; HER2-n = 32). E, F) Correlation between EVs-HER2 levels and ICH HER2 status assessed by IHC (score range from 0 to 3+) in mBC patients (CD63/HER2 p-value _(Spearman)_: n.s.; MSP/HER2 p-value _(Spearman)_: 0.03); Data are shown as box and whisker plots. Each data point represents an individual subject analysed. p-value *<0,05.

While both assays for EVs-HER2 were able to detect differences between the groups, by correlating SiMoA data with the HER2 status clinically assessed by IHC pathological assessment in mBC patients, we observed that data obtained using MSP beads demonstrated a better correlation with IHC findings than EV-HER2 levels measured using anti-CD63 conjugated beads where no meaningful correlation with HER2 status was detected (Figure 5E and 5F).

Finally, we explored the relationship between the CD9+ EVs and EVs-HER2 levels measured by the MSP assay and disease progression in mBC patients. Each mBC patient was monitored from the time of blood sample collection at the onset of metastasis until the last follow-up date (either death or transfer to another institution): the mean follow-up time for the entire mBC cohort was 17 ± 14 months. Participants were stratified into three groups based on their clinical outcomes assessed 3 years post-enrolment: deceased (n=12), disease progression (n=16), and progression-free survival (n=16). For a subset of mBC (n =5) data were not available.

The results showed that higher EVs levels at metastatic diagnosis were associated with disease-free survival (DFS), whereas lower circulating CD9+ EVs levels were linked to poorer outcomes (Figure 6A). In contrast, no significant correlation was found between EVs-HER2 levels and disease progression (Figure 6B).

**Figure 6:**
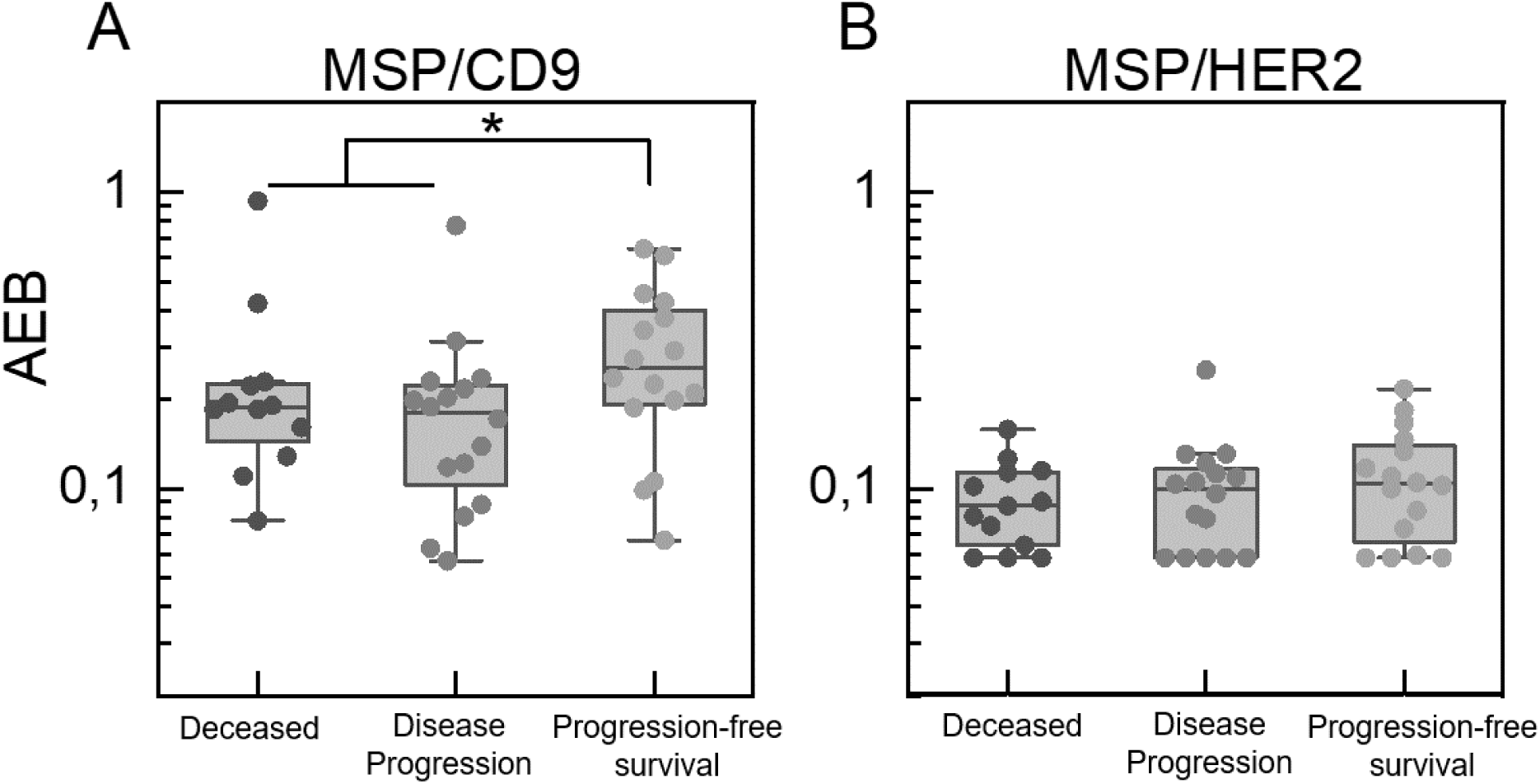
Relationship between EVs and EVs-HER2 levels measured by the MSP assay and disease progression in mBC patients. p-value *<0,05.

## 3. DISCUSSION

The translation of EVs or markers transported within or on the surface of EVs into clinical practice has long been considered a promising yet challenging approach able to provide real time information on cancer evolution in every stage of the disease^32^. Up to now, however, several obstacles have hampered their use, particularly within the oncology field, where liquid biopsy is gaining broader acceptance but EVs face fierce competition from other circulating biomarkers such as circulating tumor cells (CTCs) and circulating tumor DNA (ctDNA)^33,34^.

With the introduction of new HER2-targeted therapies into clinical practice, and emerging evidence demonstrating that even HER2-low patients may benefit from anti-HER2 treatments, the need to assess HER2 expression more sensitively and across different disease settings has become clinically relevant. Circulating HER2+ EVs might have a critical clinical relevance, as could offer a non-invasive approach to assess and monitor HER2 expression dynamically. Traditional EVs isolation and detection methods, however, struggle with low-abundance EVs, highlighting the need for sensitive, specific strategies. Challenges include stoichiometric and analytical limits, EV heterogeneity, dilution in complex biofluids, and outnumbering by other extracellular and non-vesicular particles ^11^.

This study specifically addressed the significant difficulties associated with the reliable detection and quantification of rare subpopulations of circulating EVs, as those transporting HER2, in mBC patients^35,36^, using a combination of ultrasensitive detection approach as a careful selection of binding agents.

The use of SiMoA for EVs analysis offers a scalable, easy-to-use platform for rapid, ELISA-like monitoring of multiple samples. By coupling SiMoA with the MSP capture agent, we bound EVs independently of their tetraspanin expression levels ^37,38^. To detect HER2 specifically, we employed our in-house antibody MGR2 ^23^, which recognizes a unique epitope on HER2’s extracellular domain different from the one targeted by current anti HER2 therapies thus minimizing therapeutic interference and enhancing assay specificity. An assay using MSP beads and anti-CD9 antibody was developed and used to evaluate also general EVs level in plasma.

Additionally, the two corresponding assay using magnetic beads conjugated with a commercial anti-CD63 antibody were prepared. This direct comparison of MSP-based capture versus antibody-based capture enabled us to select the most effective protocol for EV-associated HER2 analysis.

Results obtained using PDOs-derived EVs showed how while both antibodies based and MSP-based assays can be sued for the task, MSP-conjugated beads significantly enhanced assay sensitivity, allowing the detection of lower amount of EVs and a precise quantification of HER2-positive EVs. This resulted particularly evident as the MSP based assay for EVs-HER2 was able to discriminate between EVs derived from the BC-PDO and the one obtained by the HC-PDO, a task where the assay using the traditional anti-CD63 conjugated beads failed. This robust performance underscores the feasibility and translational potential of the MSP-based SiMoA assay.

Correlation of the results obtained on plasma samples revealed only modest agreement between MSP-and CD63-based assays, reflecting their differing target populations.

It is well known that, even though CD63 is considered a reliable EVs marker, not all EVs display this protein on their surface. Antibody-based bead assays thus analyse only a partial subpopulation of EVs. MSP was developed specifically to overcome this limitation, acting as pan-vesicles binders. As confirmed by our results, MSP detected a much higher number of EVs, both in PDO-derived EVs analysis, where lower EVs amounts were required for the dose-response curve, and in plasma samples, where reliablegeneralEVs quantification was achieved using a 1:100 plasma dilution— requiring only 1 µL of plasma. On the other hand, MSP sensitivity to membrane curvature decreases binding efficiency to lipid bilayers above 200 nm^37^, resulting in a greater effectiveness in quantifying small EVs. This subset may not represent all EVs and might not be universally relevant across different studies.

In accordance with MISEV guidelines and the recommendations by the ISEV Blood EV Task Force (MIBlood-EV)^31^, we evaluated the influence of pre-analytical factors on our data by spectrophotometrically assessing haemolysis and lipemia in the analysed samples. Notably, no significant correlation was observed between these pre-analytical factors and assay results, including the MSP/HER2 assay highlighting the robustness of the SiMoA EV analysis.

Concerning the analysis of EVs in mBC and HC, mBC samples surprisingly presented lower EVs levels in both antibody- and MSP-based assays. This result contrasts with our previous study on early breast cancer patients, where elevated levels of CD63+ and CD9+ EVs were detected^19^. Although the two works are not directly comparable as antibodies used in the previous work were different, this work suggest that the high EVs levels in early disease stages might not originate directly from cancer cells but represent some kind of adaptative response at the presence of the cancer. This discrepancy also raises some concerns regarding the translatability of proteomic studies conducted on EVs extracted from early breast cancer plasma samples to monitor more advanced stages of the disease^39,40^. It must be notice also that this result, although confirmed in all our assays, is in contrast with some other report on the literature where CD105+ EVs were found more elevated in women with mBC than in healthy controls^41^ and where a combination of markers on EVs were found enriched by thermophoretic aptasensor analysis in particular in patient with mBC with disease progression ^42^. Our results thus underly once more the need to consider EVs composition or multiple markers while monitoring mBC.

Although the mechanism driving the lower EV levels in mBC remains unclear, we observed consistently this result across all assays, including those for EVs-HER2; with the CD63/HER2 test failing to reach statistical significance, further confirming the lower response of this assay.

Interestingly, by the monitoring of disease progression in our patient cohort we could detect that lower EVs values measured by the MSP/CD9 assay were also associated with worse clinical outcomes, confirming the strength of the results obtained and suggesting that EVs quantification using MSP-based approaches may provide valuable prognostic information, and suggesting that the EVs levels in blood might be used to gain prognostic information. Notably the fact that also MSP showed a decrement of EVs in mBC suggest that this effect is not due only to a variation of the composition of EVs surface markers in advanced stages of the disease but that is the overall number of particles detected is decreasing. One of the possible explanation of this would be in an higher interaction of cancer released EVs with plasmatic components such as lipoproteins^43^, a phenomenon known as EVs-corona^44^, also detected in our previous work on the characterization of EVs by Raman spectroscopy^45^,

By examining differences between HER2+ and HER2-mBC patients, both assays revealed significantly higher EVs-HER2 levels in HER2+ patients, while no differences were observed in the CD9+ EVs levels. This confirms a close association between EVs-HER2 and the presence of cancer cells overexpressing HER2 on their surface^46^. It is notable to say that while the high background of the MSP/HER2 assay reduces slightly the difference of EVs-HER2 observed, in several HER2+ mBC patients the detected levels were still undetectable. The results obtained with MSP/HER2 assay correlated better with the HER2 score measured using the actual standard IHC method suggesting that this assay could provide a better way to monitor cancer evolution in mBC and that could be used to monitor the response to the treatment with anti-HER2 therapies not only in the metastatic setting but also in patient undergoing neoadjuvant treatments ^47^.

Overall, our optimized SiMoA assay employing MSP-conjugated beads and an in house developed anti-HER2 antibody targeting the extracellular domain of HER2 significantly advances the sensitive and specific detection of HER2-expressing circulating EVs. This methodological advancement holds considerable promise for clinical application in mBC, potentially facilitating accurate monitoring of HER2 status and informing timely, precise therapeutic interventions.

Some limitations of this work persist, as the assays are currently not fully optimized for being used in clinical practice, or as secondary outcome in clinical studies, as they still cannot be applied on large patient cohorts. In particular, we still lack a standardized and quality-controlled method for producing HER2-bearing EV standards, essential for achieving fully quantitative and reproducible assays. Consequently, our clinical study’s limited sample size for both mBC patients and HC restricts the clinical robustness and generalizability of our findings.

## 4. CONCLUSION

In conclusion, our study shows how SiMoA immunoassays using MSP-conjugated beads outperform traditional antibody-based tests by delivering superior sensitivity and specificity.

When used to analyse PDO derived EVs, the MSP EVs-HER2 assay unequivocally distinguished EVs from BC-PDO versus HC-PDO. When applied to clinical samples, EVs-HER2 levels were enriched in HER2-positive mBC and showed a good correlation with IHC scores, validating the assay’s biological and clinical relevance. Interestingly, CD9+ EVs levels were consistently lower in mBC patients compared to HC, and further depletion measured by the MSP assay was associated with poorer clinical outcomes suggesting that CD9+ EVs level could be of potential prognostic value. Our results thus show how MSP enhance SiMoA’s performance facilitating the monitoring of HER2 status by EVs analysis and providing information of the aggressiveness of the disease by looking at CD9+ EVs levels.

## 5. MATERIALS AND METHODS

### 5.1 EVs isolation from patient-derived organoids (PDOs)

Two patient-derived organoids (PDOs) were derived from breast cancer tissues (BC-PDO) and from healthy breast issue (HC-PDO) according to a previously published method^26^. Once the PDOs were seeded in six wells of a multiwell plate and reached confluence, the culture medium was collected every 72 hours over three time points. A total of 18 mL of PDO-conditioned medium was centrifuged at 4000 ×g for 10 minutes at 4°C to remove cellular debris. The resulting supernatant was concentrated to approximately 1 mL using Amicon Ultra-15 filters (Merck Millipore,) by centrifuging at 4000 ×g for 25 minutes at 4°C. The concentrated sample was then loaded onto a size-exclusion chromatography (SEC) column packed with 10 mL of Sepharose CL-6B resin (Merck), pre-equilibrated with phosphate-buffered saline (PBS)^48^. Fractions of 500 µL of PBS were eluted. Fraction from 1 to 6 were discarded, while fractions from 7 to 16 were collected. EVs elution start at mL 3 to mL 5. Then, the column was washed with at least 2 column volume (CV) of PBS. The EVs containing fractions (7-10) were pooled and concentrated using Amicon Ultra-2 mL filters (Merck Millipore) by centrifuging at 3200 ×g for 25 minutes at 4°C. To recover the concentrated EVs, the filter was inverted and centrifuged for 1 minute. The EVs were then stored at −80°C until further use.

### 5.2 Nanoparticle Tracking Analysis

Nanoparticle tracking analysis (NTA) was conducted to determine the size of EVs using the NS300 instrument (NanoSight, Amesbury, UK). Samples were diluted 1:100 in filtered PBS to achieve an optimal concentration of 10^8^–10^9^ particles/mL. A volume of 1 mL of the diluted sample was loaded into the instrument, and particle movement was recorded at a frame rate of 30 frames/s. For each sample, videos of particle movement were recorded three times, and the concentration and size of the EVs were determined using the NTA software (version 2.2, NanoSight).

### 5.3 Western Blotting Analysis

For Western Blotting (WB) analysis, 20 µg of the of protein (determined by BCA assay) were mixed with 4X Laemmli buffer (Biorad) to detect tetraspanins and boiled for 5 minutes at 95 °C. For the detection of Alix, the lysis of EVs was performed in RIPA buffer (50 mM Tris-HCl, 150mM NaCl, 1% Np-40, 12 mM Deoxycolic acid and 1% cOmplete Protease Inhibitor Cocktail) for 20 minutes in ice. Following centrifugation at 18,000 × g for 20 minutes at 4 °C, the protein content was measured using the BCA assay. Sample were resuspended in reducing Laemmli buffer supplemented with β-mercaptoethanol as a reducing agent and boiled for 5 minutes at 95°C.

Proteins were separated on 10-12% SDS-PAGE polyacrylamide gels and transferred to Polyvinylidene fluoride (PVDF) membranes (Bio-Rad Laboratories). The blocking step was carried out with Tris-buffered saline containing 0.1% Tween (TBS-T), supplemented with 5% bovine serum albumin or 5% fat-milk for 1 hour at room temperature. Membranes were incubated overnight at 4°C with the following antibodies diluted in blocking buffer: anti-CD63 (1:1000, HansaBiomed), anti-CD81 (1:1000, Ancell), anti-Alix (1:1000, Santa Cruz Biotechnology). The membranes were washed three times with TBS-T, repeated twice, and then incubated with the secondary anti-rabbit antibody or the secondary anti-mouse antibody conjugated with horseradish peroxidase (both antibodies diluted 1:5000 in blocking buffer, Abcam) for 2 hours at room temperature. The membranes were washed three times with TBS-T and analysed with a ChemiDoc System (Bio-Rad Laboratories).

### 5.4 anti CD63 based SiMoA Assays

Following the manufacturer’s guideline, the paramagnetic carboxylated beads (150 μL) (Quanterix) were activated with 1-ethyl-3-(3-dimethylaminopropyl) carbodiimide hydrochloride (EDC, Sigma-Aldrich), subsequently conjugated with the anti-CD63 capture antibody (HansaBiomed) and stored in the Bead Diluent buffer (Quanterix) at 4 °C until use.

Isolated EVs were diluted appropriately in Homebrew Sample Diluent (Quanterix), with a final volume of 100 µL. Additionally, 25 µL of paramagnetic beads, previously conjugated, were added. Next, 20 µL of anti-CD9-biotin (Ancell) or anti-HER2 MGR2 antibody —biotinylated prior to use according to the manufacturer’s instructions using Pierce EZ-Link NHS-PEG4 (Thermo Fisher)— were added to each well at a working concentration of 1 µg/mL.-. The mixture was incubated for 30 mi nutes at 25 °C with continuous shaking at 800 rpm. After incubation, beads are washed with an automatic plate-washer and then each incubated for 10 min with a 100 µL of Streptavidin-ß-galactosidase (in SBG Diluent, Quanterix). After SBG incubation step, the plate is washed again by the automatic plate-washer and then inserted into the Quanterix SR-X instrument for analysis where Resorufin ß-D-galactopyranoside (RGP, Quanterix) was automatically added. Detection and quantification were performed autonomously by the system.

### 5.5 Membrane Sensing Peptides (MSP) based SiMoA Assays

Following the manufacturer’s guideline, 150 μL of carboxylated paramagnetic beads from the Quanterix Homebrew assay SiMoA kit (2.8 × 10^9^ prt/ml) were activated with EDC, then 300 μL NH_2_-PEG4-N_3_ (from Sigma-Aldrich) solution 1 mM in PBS was added and shaken for 2 hours. After washing twice with Bead Wash Buffer (Quanterix) to remove excess reagent, the beads were incubated with 300 μL of 100 µM MSP-DBCO solution in PBS for 1 hour under mixing. After the conjugation step, the beads were washed two times with Bead Wash Buffer (Quanterix) and then were blocked with Bead Block Buffer (Quanterix) for 15 min at room temperature under shaking and subsequently washed with Bead Wash Buffer (Quanterix). The beads were stored in Bead Diluent (Quanterix) at 4 °C until use.

Isolated EVs were diluted appropriately in Homebrew Sample Diluent (Quanterix), with a final volume of 100 µL. For each well 25 µL of paramagnetic beads, previously conjugated, were added into a 96-well microplate. The mixture was incubated for 2 hours at 30 °C with continuous shaking at 800 rpm. Following the incubation, the beads were washed using an automatic plate washer, then incubated with anti-CD9-biotin antibody or anti-HER2-biotin MGR2 antibody for 10 minutes. After incubation, beads were washed with an automatic plate-washer and then each incubated for 10 min with a 100 µL of SBG (in SBG Diluent, Quanterix). After a washing step, the plate and RGP substrate (Quanterix), were loaded into a Quanterix SR-X instrument for analysis. Detection and quantification were performed autonomously by the system.

### 5.6 Patient selection

Forty-nine mBC patients and thirty HC, were enrolled in the protocol NCT05798338 approved by the Ethical Committee of ICS Maugeri IRCCS (Pavia, Italy) and conducted in compliance with the Declaration of Helsinki. All patients who agreed to participate signed a specific informed consent prior to the inclusion in the study. The main clinical characteristics of the study cohorts are summarized in Table 2.

HER2 status in mBC was evaluated according to current American Society of Clinical Oncology (ASCO) and the College of American Pathologists (CAP) guidelines, based on IHC and Fluorescence in situ hybridization (FISH) analysis^3^. Statistical analysis: t-Test. p-value *<0,05. n.s, not significant.

### 5.7 Blood collection

Blood samples were collected in EDTA-coated tubes. The blood samples were then centrifuged at 2000×g for 10 minutes at room temperature. Plasma samples were collected and centrifuged a second time at 2500×g for 10 minutes at 4 °C. The plasma was then stored at −80 °C until use, by the Institutional Biobank “Bruno Boerci”.

### 5.8 EVs detection using the anti-CD63 based SiMoA assays

The protocol outlined in Section 2.4 was applied for the analysis of human plasma samples. Plasma samples were diluted 1:4 in Homebrew Sample Diluent (Quanterix) and processed according to the established procedure. Immunoassay analysis was performed using anti-CD9-biotin or anti-HER2-biotin antibodies in combination with anti-CD63 magnetic beads.

### 5.9 EVs detection using the MSP based SiMoA assays

The protocol outlined in Section 2.5 was applied for the analysis of human plasma samples. Plasma samples were diluted 1:4 in Homebrew Sample Diluent (Quanterix), for the HER2 assay and 1:100 for the CD9 assay, following the established procedure. Immunoassay analysis was performed using anti-CD9-biotin or anti-HER2-biotin antibodies in combination with MSP-conjugated beads.

### 5.10 Statistical Analysis

For the statistical analysis, Kolmogorov-Smirnov and Shapiro-Wilk tests were applied to verify the normal distribution of data. Then, parametric [t test] or non-parametric [Mann-Whitney test] tests were applied as appropriate to compare mean values between the different groups using Origin software. Statistical significance was set at p-value < 0,05. Correlation between preanalytical factors (Table S1) and EVs values were performed by Spearman analysis.

## ACKNOWLEDGEMENTS

This work was supported by the Italian Ministry of University and Research under the scheme “PRIN: progetti di ricercar di rilevante interesse nazionale– Bando 2022, Prot. 2022CS9H53” and by the Italian Ministry of Health “Ricerca corrente 2025”.

We want to thank the Institutional Biobank “*Bruno Boerci*” for the support in the storage and handling of the clinical samples analysed. We extend our deepest gratitude to all the patients who participated in this study. We thank Raffaele Allevi, for assistance with transmission electron microscopy (TEM) imaging. This work is dedicated to the memory of Dr. Marina Cretich that died for cancer about one year ago and that was the original principal investigator of the EV-Print project that supported with work. She made a decisive contribution to this work and that continue to inspire us.

The Graphical abstract was created in BioRender.

## DATA AVAILABILITY STATEMENT

The datasets used and analysed during the current study are available from the corresponding author upon reasonable request.

## ETHICS APPROVAL STATEMENT

This study was approved by the Ethical Committee of ICS Maugeri (Pavia, Italy) authorized the study as protocol NCT05798338.

## DECLARATION OF INTEREST STATEMENT

A.G. is listed as co-inventor on European patent application EP4025911A1 (Conjugates composed of membrane-targeting peptides for extracellular vesicles isolation, analysis and their integration thereof). The patent covers the peptide-based probes and methods described in this study. All other authors declare no competing interests.

## DECLARATION OF GENERATIVE AI AND AI-ASSISTED TECHNOLOGIES IN THE WRITING PROCESS

During the preparation of this work the author(s) used ChatGTP in order to revise English language and style. After using this tool/service, the author(s) reviewed and edited the content as needed and take(s) full responsibility for the content of the publication.

## Notes

### Competing Interest Statement

The authors declare the following potential conflicts of interest:
Alessandro Gori (A.G) is listed as co-inventor on European patent application EP4025911A1 (Conjugates composed of membrane-targeting peptides for extracellular vesicles isolation, analysis and their integration thereof). The patent covers the peptide-based probes and methods described in this study. All other authors declare no competing interests.

